# PRMT5 promotes symmetric dimethylation of RNA processing proteins and modulates activated T cell alternative splicing and Ca^2+^/NFAT signaling

**DOI:** 10.1101/2021.08.24.457384

**Authors:** Shouvonik Sengupta, Kelsi O. West, Shridhar Sanghvi, Georgios Laliotis, Laura M. Agosto, Kristen Lynch, Philip Tsichlis, Harpreet Singh, Kristin Patrick, Mireia Guerau-de-Arellano

## Abstract

Protein Arginine Methyltransferase (PRMT) 5 is the major type 2 methyltransferase catalyzing symmetric dimethylation (SDM) of arginine. PRMT5 inhibition or deletion in CD4 Th cells reduces TcR engagement-induced IL-2 production and Th cell expansion and confers protection against experimental autoimmune encephalomyelitis (EAE), the animal model of Multiple Sclerosis. However, the mechanisms by which PRMT5 modulates T helper (Th) cell proliferation are still not completely understood and neither are the methylation targets in T cells. In this manuscript, we uncover the role of PRMT5 on alternative splicing (AS) in activated T cells and identify several targets of PRMT5 SDM involved in splicing. In addition, we find a possible link between PRMT5 mediated AS of *Trpm4* (Transient Receptor Potential Cation Channel Subfamily M Member 4) and TcR/NFAT signaling/IL-2 production. This understanding may guide development of drugs targeting these processes to benefit patients with T cell-mediated diseases.

## Introduction

CD4 T helper (Th) cells arguably play one of the most critical roles in immunity, by orchestrating antigen-specific adaptive immunity and enhancing innate immunity via release of cytokines^1^. The resulting cytokine gradient elicits autocrine and paracrine effects on CD4 Th cells, CD8 T cytotoxic cells, B cells and myeloid cells. Therefore, a lack of CD4 Th cells substantially impacts both humoral and cytotoxic immune responses and commonly results in life-threatening infections. In turn, over-reactive CD4 Th cell responses can lead to the chronic inflammation and tissue destruction observed in autoimmune disease. Protein Arginine Methyltransferase (PRMT) 5 is a Type II methyltransferase that catalyzes symmetric dimethylation (SDM) of protein arginines and plays an important role in development and cancer. Previous work from our lab and others has shown that PRMT5 is induced after CD4 Th cell activation/autoimmune responses, and that loss of protein arginine methyltransferase (PRMT)5 reduces TcR engagement-induced Th cell expansion and confers protection against the mouse model of Multiple Sclerosis, experimental autoimmune encephalomyelitis (EAE)^2–4^. However, the methylation targets of PRMT5 in T cells and associated molecular mechanisms are not well defined^5^.

A key step for protective immune or pathogenic autoimmune responses is the clonal expansion of antigen-specific T cells induced by TcR engagement^6^. TcR engagement activates signaling pathways^7^ that lead to Nuclear factor of activated T-cells (NFAT) activation^8^ and cell cycle progression^9^. NFAT activation results in nuclear localization, activation of the IL-2 promoter and IL-2 cytokine transcription^10^. Once secreted, IL-2 binds the IL-2 receptor in an autocrine and paracrine manner and promotes T cell growth and proliferation^11^. We have previously seen that PRMT5 can promote IL-2 production, cell cycle progression^12^ and T cell proliferation^2^. However, the impact of PRMT5 loss on TcR/NFAT signaling leading to IL-2 production and T cell proliferation remains unexplored.

As a consequence of TcR signaling, T cells undergo dramatic changes in their gene expression programs. These changes support the transition from naïve to highly proliferating and cytokine-producing effector T cells. A substantial portion of gene expression modulation occurs at the gene expression level. However, additional modulation is possible via alternative splicing (AS)^13^. AS is the process by which exons are included or excluded in the final processed mRNA transcript, resulting in distinct isoforms from the same gene^14^. AS therefore provides an important layer of gene expression programming control, by diversifying the proteins that are actually encoded within genes. Previous work from the Lynch lab has established that antigenic/TcR stimulation modulates the AS gene expression pattern of T cells^13,15,16^ as revealed by RNA-Seq, quantitative microarray, bioinformatics and RT-PCR analyses. The resulting protein isoforms have been linked to functional outcomes such as TcR α chain transcription^17^, TcR signal transduction^18^ and JNK–CELF2 dependent splicing control^19^, indicating AS plays crucial functional roles in activated T cell biology.

In this manuscript, we explore the specific role of PRMT5 on AS changes induced after T cell activation, methylation targets of PRMT5 in T cells involved in splicing and potential links between a specific AS *Trpm4* isoform and altered TcR/NFAT signaling. We find that PRMT5 deletion alters the AS pattern induced by T cell activation and results in the loss of SDM of proteins involved in splicing, such as SmD and hnRNPK. We also report specific validated changes in the AS of *Trpm4*, a Ca^2+^ responsive Na+ channel that plays an important role in total calcium processing and NFAT dependent IL-2 production in Th cells. Overall, these data conclusively link PRMT5 to TcR-induced AS in T cells and suggest that altered methylation in splicing proteins and changes in Ca^2+^/NFAT signaling underlie TcR expansion defects in PRMT5 deficient T cells.

## Materials and Methods

### Mice

Age-matched 9-13 week-old iCD4-PRMT5^fl/fl^ (CD4creER^−^PRMT5^fl/fl^) and iCD4-PRMT5^Δ/Δ^ (CD4creER^+^PRMT5^fl/fl^) mice, described in Webb *et al*^3^, on the C57BL/6 background were used for RNA-SEq and mass spectrometry. Age-matched 9-13 week-old C57BL/6 background constitutive T-PRMT5^fl/fl^ (CD4cre^−^PRMT5^fl/fl^) and T-PRMT5^Δ/Δ^ (CD4cre^+^PRMT5^fl/fl^) mice, also described in Webb *et al*^3^, were used in the remainder of experiments. Males and females were used in experiments and no significant differences were observed between genders. Animal use procedures were approved under Institutional Animal Care and Use Committee protocol number 2013A00000151-R1. All animals were euthanized under the American Veterinary Medical Association (AVMA) guidelines.

### Deletion of PRMT5 and murine CD4 Th cell isolation *in vivo* tamoxifen treatment

iCD4-PRMT5^fl/fl^ and iCD4-PRMT5^Δ/Δ^ mice were administered 300mg/kg (7.5 μL/g) tamoxifen (Sigma-Aldrich, catalog no. T5648) by gavage for 5 days, and euthanized 2 days after the last dose for secondary lymphoid organ (lymph nodes and spleen) harvest. Deletion of PRMT5 in T cells did not require tamoxifen treatment in T-PRMT5^fl/fl^ (CD4cre^−^PRMT5^fl/fl^) and T-PRMT5^Δ/Δ^ (CD4cre^+^PRMT5^fl/fl^) mice. Harvested organs were processed to a cell suspension and subsequently used for CD4 Th cell isolation. Murine CD4 T cells were isolated from processed lymphoid organs using EasyEights magnet (Stem Cell Technologies, Catalog no. 18103) and the whole CD4^+^ T cell isolation kit (Stem Cell Technologies, catalog no. 19852). Purity of CD4 T cells was in the range of 87-95%, as measured by flow cytometry. Additional details on the tamoxifen treatment regimen and mouse immunological parameters after tamoxifen treatment can be found in Webb *et al*^3^.

### Cell culture

T cells were cultured in EAE media (RPMI + 10% FBS + 2mM L-glutamine + 1:100 Penn-Strep + 1mM Sodium Pyruvate + 1:100 Minimal essential amino acids + 13mM HEPES + 1:500 β-mercaptoethanol). Unless otherwise indicated, CD4 T cells were activated on coated 5 μg/mL CD3 and soluble 2 μg/mL CD28 for 48hr in 24-well plates. Human Jurkat T cells with a stable *PRMT5* knockdown were generated by the Tsichlis lab at OSU, as previously described^3^. Briefly, pLx304 DEST EV was used as an empty vector control cell line (termed EV) and PRMT5 shRNA (MilliporeSigma, catalog no. SHCLNG-NM_006109, clone ID TRCN0000107085) was used to induce the PRMT5 knockdown cell line (termed shPRMT5). Cells were cultured in standard Jurkat cell culture media (ATCC) for 24-48hr until desired confluency was reached prior to downstream processing.

### RNA-Seq

Whole CD4^+^ T cells from iCD4-PRMT5^fl/fl^ and iCD4-PRMT5^Δ/Δ^ mice (n = 3 pooled mice/sample and n = 3 samples per group) were used for RNA-Seq. Samples were either lysed directly *ex vivo* (resting) or activated (anti-CD3/CD28, no cytokines, 48 hr) before lysis and RNA isolation. RNA isolation was done with the Direct-zol RNA Miniprep (Zymo Research, catalog no. R2052) according to the manufacturer’s instructions. 1 ng of total RNA was used for quality control (QC), library preparation, and RNA-Seq. Quality of RNA was evaluated using the Agilent 2100 Bioanalyzer and RNA Nano chip (Agilent Technologies). Samples with RNA integrity number (RIN) greater than 7.7 were considered for sequencing. Files pertaining to activated T cells associated with the RNA-Seq experiment can be found in NCBI’s Gene Expression Omnibus (GEO) under the accession number GSE141168. For additional information on RNA-Seq run and analysis, refer to protocol listed in Webb *et al*^3^. RNA-Seq was performed by the Genomic Services Laboratory of the Abigail Wexner Research Institute at Nationwide Children’s Hospital, Columbus, Ohio.

### MAJIQ and VOILA

Alternative splicing events were analyzed using MAJIQ and VOILA under default parameters (Vaquero-Garcia et al., 2016). PRMT5^fl/fl^ mice T cell FastQ files were set as the control group to compare PRMT5^Δ/Δ^ files against. In brief, raw junction spanning reads from RNA-Seq fastQ files were aligned to the GRCm38.p3 assembly of the *Mus musculus* reference from NCBI using STAR RNA-Seq aligner (2.6.0c). These alignments were fed into MAJIQ to construct splice graphs for transcripts using the RefSeq annotation and identify both known and novel alternative splicing events in the dataset. All identifiable local splice variants (LSVs) were analyzed from the splice graphs with minimum reads set to at least 10 to pass the quantifiable threshold. For each exonic-intronic junction in an LSV, MAJIQ quantified the expected percent spliced (Ψ) value in PRMT5^fl/fl^ and iCD4-PRMT5^Δ/Δ^ T cell samples and the expected change in Ψ (ΔΨ) between PRMT5^fl/fl^ and iCD4-PRMT5^Δ/Δ^ T cell samples. The VOILA results were processed with a filter of at least 20% to include high confidence changing LSVs (at least two junctions with a 95% probability of expected ΔΨ of at least an absolute value of 20 Ψ units (ΔΨ ≥/≤ 20) between PRMT5^fl/fl^ and iCD4-PRMT5^Δ/Δ^ T cell samples. The high confidence results were further classified into exon skipping, alternative 5’, alternative 3’ splice site or intron-retention events.

### Semi quantitative PCR

To evaluate mRNA expression, 200–300 ng of RNA were reverse transcribed using oligo d(T) or random primers and Superscript III (Thermo Fisher Scientific, catalog no. 18080051) according to the manufacturer’s instructions. Samples were run on a Nexus mastercycler (Eppendorf). Exon 20 region specific primers spanned from exon 19 to exon 21 (Fwd: TCCTCTTCTTCCTCTGCGTG, Rev: ATTCCCGGATGAGGCTGTAG. Products-e20 skipped band - 230bp, e20 included band – 408bp). Control primers were on exon 19 (Fwd: CCTCTTCTTCCTCTGCGTGT, Rev: ATTTCCTCCTGGGGAATTTG. Product – 150bp). An initial denaturation step at 95°C for 5 minutes was followed by 30 cycles of denaturation at 95°C for 30s, annealing at 54°C for 1 min, extension at 72°C for 30s. PCR products were run on 1.5% agarose gels with 0.5% TBE buffer. E20 skipped PCR products were confirmed by use of nested primers (Fwd: GCC CTC ATG ATT CCA GGT AA, Rev: TCC AGT AGA GGT CGC TGT TG) and Sanger sequencing was performed at the OSUCCC genomic shared resources.

### Assessment of calcium signaling in T cells

Isolated activated (anti-CD3/CD28 – 2.5ug/ml, 50U IL-2, 48hr) CD4^+^ T cells from PRMT5^fl/fl^ and T-PRMT5^Δ/Δ^ mice were plated on poly-L lysine (Millipore Sigma, catalog no. P8920-100ML) coated glass-bottom dishes (Cellvis 35 mm - 14 mm micro-well #1.5 cover glass, Fisher Scientific, catalog no. NC0794151) for 120 min. Cells were then treated with 10 μmol Fluo-4-AM (Invitrogen, catalog no. F14201) dye for 30 min in DMEM (without phenol red and glutamine; catalog no. 11054020) at 5% CO_2_ in a humidifying incubator at 37°C. Then the dye was washed out and cells were incubated for 30 min in modified EAE media supplemented with 10% FBS for de-esterification. Following de-esterification, cells were switched to modified Ringer’s solutions with 0 mM Ca^2+^ (120 mM NaCl, 5 mM KCl, 1 mM MgCl_2_, 25 mM NaHCO_3_, and 5.5 mM D-glucose, pH 7.3) for imaging with a Nikon A1R-HD laser-scanning confocal microscope. Fluo-4 was excited using 488 nm laser and fluorescence emission was detected at 500-550 nm. Resting Ca^2+^ baseline was recorded for 150 sec prior to addition of sarcoplasmic reticulum Ca^2+^-ATPase (SERCA) inhibitor thapsigargin (2μM). After 150 sec CaCl_2_ (2mM) was added and calcium uptake was monitored for 600 sec. The data are represented as ΔF/F_0_ vs. time, where F_0_ is basal fluorescence and ΔF=F-F_0_.

### Immunocytochemistry

Isolated activated (5 μg/ml anti-CD3 and soluble 2 μg/ml CD28, 48hr) CD4^+^ T cells from PRMT5^fl/fl^ and T-PRMT5^Δ/Δ^ mice were plated on poly-L lysine (Millipore Sigma, catalog no. P8920-100ML) coated glass cover slips for 120min. Cells were then stained with wheat germ agglutinin for 10 min prior to fixing with 4% paraformaldehyde (Electron Microscopy Sciences, catalog no. 15713) for 10 min and permeabilization with 0.5% Triton-X 100 for 10 min. Samples were blocked with 10% normal goat serum for 10 min and incubated in NFATc1 antibody (Santa Cruz Biotechnology, catalog no. sc-7294) overnight at 4°C. Samples were then incubated in secondary antibody conjugates Atto 647N (1 μg/mL each of anti-mouse; Sigma catalog no. 50185-1ML-F) for 60min, followed by 10.9 mM DAPI (1:10,000) (Sigma catalog no. D9542) staining for 10 min. Coverslips were mounted with ProLong™ gold antifade (Invitrogen, catalog no. P36930) and cells were imaged with Nikon A1R high-resolution confocal microscopy. NFAT and nuclear stain colocalization index^20^ was calculated using ImageJ. Pearson’s R value with no threshold condition was selected for the calculation of colocalization index.

### Flow cytometry

For flow cytometry, 48-hour activated T cells from PRMT5^fl/fl^ and T-PRMT5^Δ/Δ^ mice were fixed with 4% paraformaldehyde (Electron Microscopy Sciences, catalog no. 15713) for 10 min in V-bottom plates (Costar, catalog no. 3897). Samples were blocked with 5% normal goat serum for 1 hr and incubated in anti-TRPM4 (Abcam, catalog no. ab106200) or normal mouse IgG (Santa Cruz Biotechnology, catalog no.sc-2025) antibodies overnight. Samples were then incubated in goat anti-rabbit IgG Alexa Fluor-488 conjugated (Abcam, catalog no. ab150085) secondary for 60 min the next day prior to washing and running on FACSCalibur with DxP multicolor upgrades (Cytek). Analysis was performed using FlowJo_V10.

### Cytokine ELISA

IL-2 cytokine in 48 hr supernatants of activated T cells from PRMT5^fl/fl^ and T-PRMT5^Δ/Δ^ mice was analyzed by sandwich enzyme-linked immunosorbent assay (ELISA). Murine IL-2 ELISA capture (catalog no. 14-7022-85) and detection (catalog no. 13-7021-85) antibody reagents were purchased from Invitrogen/eBioscience. The capture antibody was coated overnight at 2 μg/ml in coating buffer (0.1M NaHCO_3_, pH 9.5). The following day, plates were washed with 0.1% PBS/Tween-20 solution and blocked with 1% BSA/PBS for 2 hr. Following blocking, 100 μL of IL-2 standard (Invitrogen eBioscience, catalog no. 14-8021-64) or supernatants are added to the wells. Plates were incubated overnight at 4°C and, the following day, plates were washed with 0.1% PBS/Tween-20 solution and 100 ul of detection antibody diluted in 1% BSA/PBS was added to the wells for 60 min, followed by 2.5 μg/ml avidin-peroxidase prepared in 1% BSA/PBS for 30 min. After washes, 0.1% H_2_O_2_/ABTS was added to the wells and the developed color signal was read at 405 nm on the SpectraMax Plus 384 plate reader (Molecular devices) at 2 – 15 min.

### Mass spectrometry

Isolated resting and activated (anti-CD3/CD28, no cytokines, 2 days) CD4^+^ T cells from iCD4-PRMT5^fl/fl^ and iCD4-PRMT5^Δ/Δ^ mice (*n* = 3 pooled mice/sample and *n* = 3 samples/group) were lysed in our in-house lysis buffer (50 mM triethylammonium bicarbonate, Millipore Sigma, catalog no. T7408-500ML + 0.05% n-Dodecyl β-D-maltoside, Millipore Sigma, catalog no. D4641-1G). Protein was quantified using Pierce BCA kit (Thermo Fisher, catalog no.23225) and 30 μg was used for immunoprecipitation (IP). IP with the SYM10 antibody (Millipore Sigma, catalog no.07-412) was done according to manufacturer’s instructions using the Pierce A/G magnetic beads (Thermo Fisher, catalog no.88802). Liquid Chromatography with tandem mass spectrometry (LC-MS/MS) was performed on IP samples using an Orbitrap fusion mass spectrometer equipped with an EASY-Spray™ Sources operated in positive ion mode by the OSU-Genomics Shared Resources (GSR). Samples were separated on an easy spray nano column (Pepmap^TM^ RSLC, C18 3μ 100A, 75μm X150mm Thermo Scientific) using a 2D RSLC HPLC system from Thermo Scientific. Each sample was injected into the μ-Precolumn Cartridge (Thermo Scientific,) and desalted with 0.1% Formic Acid in water for 5 minutes. The injector port was then switched to inject the sample and the peptides were eluted off of the trap onto the column. Mobile phase A was 0.1% Formic Acid in water and acetonitrile (with 0.1% formic acid) was used as mobile phase B. Flow rate was set at 300nL/min Mobile phase A was 0.1% Formic Acid in water and acetonitrile (with 0.1% formic acid) was used as mobile phase B. Flow rate was set at 300nL/min. Typically, mobile phase B was increased from 2% to 35% to 55% in 125 and 23 min and then increased from 55 to 90% in 10min and then kept at 95% for another 5 min before being brought back quickly to 2% in 2 min. The column was equilibrated at 2% of mobile phase B (or 98% A) for 15 min before the next sample injection. MS/MS data was acquired with a spray voltage of 1.7 KV and a capillary temperature of 275 °C is used. The scan sequence of the mass spectrometer was based on the preview mode data dependent TopSpeed™ method: the analysis was programmed for a full scan recorded between *m/z* 375 – 1700 and a MS/MS scan to generate product ion spectra to determine amino acid sequence in consecutive scans starting from the most abundant peaks in the spectrum in the next 3 seconds. To achieve high mass accuracy MS determination, the full scan was performed at FT mode and the resolution was set at 120,000. EASY-IC was used for internal mass calibration. The AGC Target ion number for FT full scan was set at 4 x 10^5^ ions, maximum ion injection time was set at 50 ms and micro scan number was set at 1. MSn was performed using ion trap mode to ensure the highest signal intensity of MSn spectra using both HCD methods (30%). The AGC Target ion number for ion trap MSn scan was set at 10000 ions, maximum ion injection time was set at 30 ms and micro scan number was set at 1. Dynamic exclusion is enabled with a repeat count of 1 within 60s and a low mass width and high mass width of 10ppm.

### Mass spectrometry analyses

Label free quantitation^21^ was performed using the spectral count approach, in which the relative protein quantitation is measured by comparing the number of MS/MS spectra identified from the same protein in each of the multiple LC/MSMS datasets. Scaffold (Proteome Software, Portland, OR) was used for data analysis. Results were filtered with 95% confident level first. Only proteins pass 1% FDR and have a minimal of 2 unique peptides were considered as valid identification.

### Western blotting and immunoprecipitation

Activated whole CD4^+^ T cells and Jurkat cells were pelleted and frozen at −80°C. Samples were lysed in RIPA buffer (10 mM Tris pH 7.4, 150 mM NaCl, 1% Triton X-100, 0.1% SDS, 1% deoxycholate) for western blotting (WB) and IP lysis buffer for immunoprecipitation (50mM Tris, 150mM NaCl, 1% NP-40, 0.1% Triton-X 100, 0.5% Sodium Deoxycholate). 4-10 μg of protein was run for the WB and 40-50 μg of initial protein was used for IP. Input samples were loaded as 10% of IP protein loading. Samples were run on 7.5% SDS-PAGE gels and transferred onto nitrocellulose membranes. Blots were blocked with 1%milk protein in TBS-Tween(0.1%). IP was performed according to manufacturer’s instructions provided by Santa Cruz Biotechnology. IP antibodies used were hnRNP K (Abcam, catalog no. - ab39975) and normal mouse IgG (Santa Cruz Biotechnology, catalog no. sc-2025). Protein A/G Plus Agarose beads were used for the pull down (Santa Cruz Biotechnology, catalog no. sc-2003). Additional information on protein isolation, western blotting, IP and blot imaging procedures can be found in Webb *et al*^3^.

### Statistics

Statistical analyses were done using the GraphPad Prism software (v9). 2-tailed Student’s t test or One-way ANOVA followed by Tukey’s or Sidak’s post hoc multiple-comparisons test was performed as appropriate. Raw RNA-Seq data was normalized and post-alignment statistical analyses were performed using DESeq2 and custom analysis scripts written in R.

## Results

### PRMT5 modulates T cell activation-dependent alternative splicing

Substantial AS modulation occurs in response to TcR stimulation of Jurkat T cells^15^. However, the contribution of PRMT5 to TcR stimulation-dependent AS changes is unknown. To address this, we leveraged our recently developed conditional CD4 T cell-specific PRMT5knockout (KO) mouse model^3^, RNA-Seq and bioinformatics tools to identify and analyze alternative splicing events. We isolated purified CD4 Th cells from iCD4-PRMT5^fl/fl^ and iCD4-PRMT5^Δ/Δ^ mice in resting vs. anti-CD3/CD28-activated conditions (henceforth referred to as activated/abbreviated as act) for paired-end RNA-Seq (**Fig. 1A**). Detection, quantification and visualization of local splicing variants (LSV) from RNA-Seq data was then achieved with the MAJIQ (Modeling Alternative Junction Inclusion Quantification) and Voila software packages created in the Barash lab^22^. MAJIQ software is, in theory, capable of detecting any splicing events involving two or more junctions, including not previously annotated splicing events.

**Figure 1.**
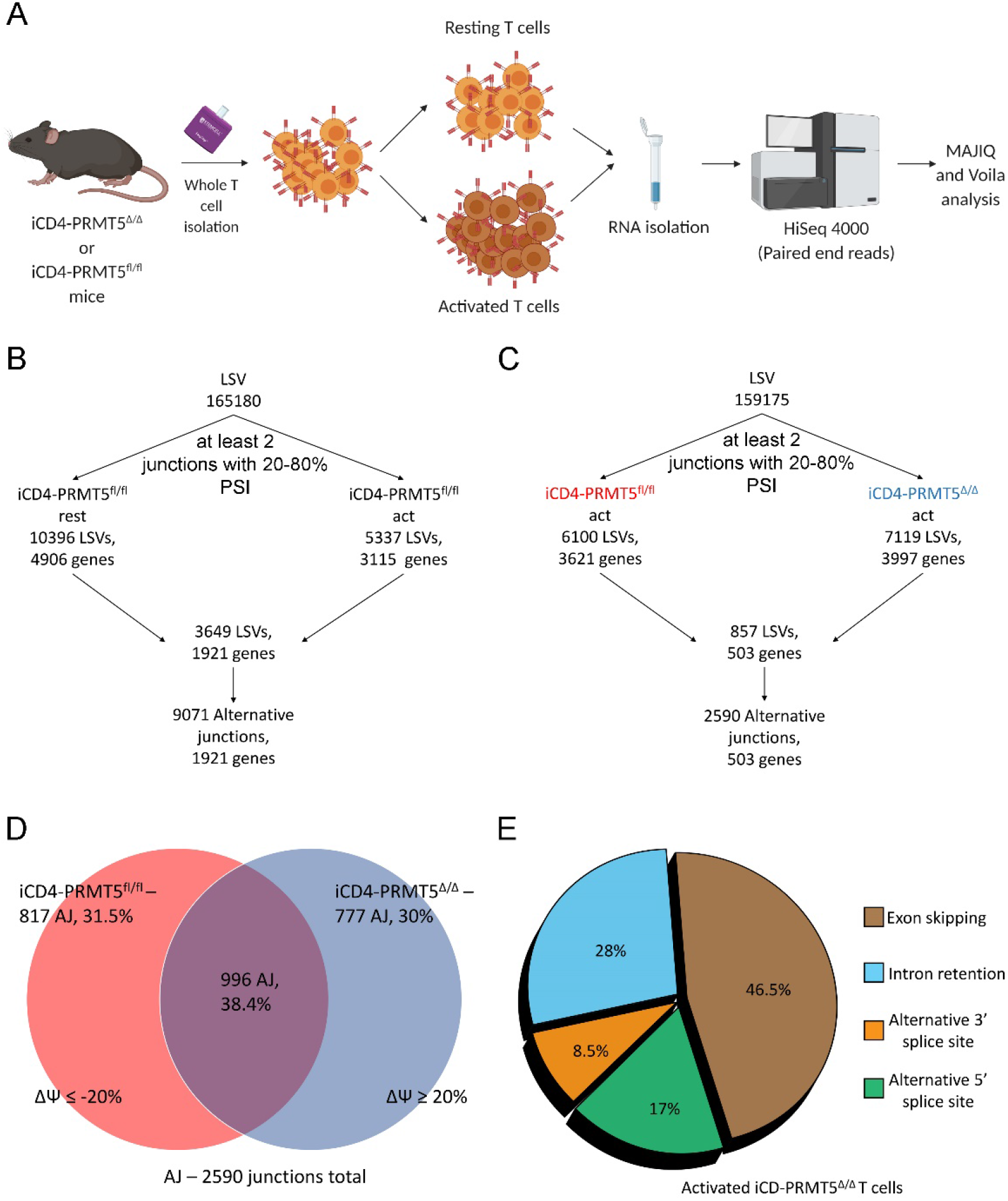
MAJIQ analysis reveals novel PRMT5 dependent changes in the alternative splicing of 503 genes in T cells. (**A**) Experimental design for paired end RNA sequencing (RNA-Seq) of resting and 48hr activated, whole CD4^+^ T cells isolated from iCD4-PRMT5^fl/fl^ and iCD4-PRMT5^Δ/Δ^ mice. n=3 samples, combined from 9 age-matched mice in each respective group. RNA-Seq was performed on the HiSeq4000 platform with approximately 60 - 80 million reads. Figure created in Biorender. (**B,C**) Analysis of local splice variants (LSVs) and alternative junctions (AJs) from MAJIQ. Workflow identifying significantly utilized AJs by comparing LSVs in resting (**B**) or 48hr activated (act) (**C**) iCD4-PRMT5^fl/fl^ or iCD4-PRMT5^Δ/Δ^ CD4^+^ T cells. LSVs with at least two or more exon junctions within the 20-80% percent spliced index (Ψ) of reads were calculated. 3649 LSVs were observed for the wild-type (iCD4-PRMT5^fl/fl)^ resting vs activated comparison (**B**) and 857 LSVs were observed for the iCD4-PRMT5^fl/fl^ act (**red**) vs iCD4-PRMT5^Δ/Δ^ act (**blue**) comparison (**C**). (**D**) Allocation of LSVs with two or more AJs in iCD4-PRMT5^fl/fl^ act (**red**) and iCD4-PRMT5^Δ/Δ^ act (**blue**) groups. Shift in AJ usage is denoted as the difference in Ψ (ΔΨ) and is set at a minimum of 20% between conditions. 817 AJs are PRMT5^fl/fl^ specific and 777 AJs are PRMT5^Δ/Δ^ specific. (**E**) Classification of 503 genes in the iCD4-PRMT5^Δ/Δ^ act group from MAJIQ shows most of the alternative splicing (AS) events belong to exon skipping (ES). Intron retention (IR) is the next largest portion of alternative splicing changes.

We first used MAJIQ to evaluate AS changes occurring as a consequence of primary murine CD4 Th cell activation in control, PRMT5-sufficient T cells. We tuned our analysis to identify the LSVs (defined as all possible splits in the exon boundary that have events), alternative junctions (AJs, defined as LSVs with two or more junctions having 20-80% of reads coming into/going out of the junction being considered) and the genes where these AS changes occurred. MAJIQ identified 9071 AJs corresponding to 3649 events impacting 1921 genes (**Fig. 1B**) upon activation of primary CD4 Th cells (comparing resting vs activated PRMT5^fl/fl^ T cells). Next, we evaluated whether the alternative splicing pattern observed in activated T cells was altered by PRMT5 deletion (comparing activated PRMT5^fl/fl^ vs PRMT5^Δ/Δ^ T cells). We observed that PRMT5 loss resulted in changes in 2590 AJs corresponding to 857 splicing events over 503 genes (**Fig. 1C**). These results indicate that PRMT5 regulates an important portion (~16%) of the activated T cell AS gene expression profile.

To determine whether there were activated T cell AJs that were unique to PRMT5 deficient or PRMT5-sufficient conditions, we performed a ΔΨ (change in percent spliced index) analysis on AJs showing splicing site usage 20% of the time or higher. Doing so, we see almost an even split of alternative junctions present in each group – unchanged (below 20% splicing site usage: 38.4% or 996 AJs), PRMT5 sufficient (ΔΨ≤-20%: 31.5% or 817 AJs) and PRMT5 deficient (ΔΨ≥20% - 30% or 777 AJs) (**Fig. 1D**). Similar analyses comparing PRMT5 sufficient vs deficient T cells in the resting state showed limited AS occurring in resting T cells, with few, albeit some differences in AS splicing (408 AJs corresponding to 127 LSVs over 74 genes, Supplemental Figure 1). Collectively, the above data indicate that the loss of PRMT5 leads to a distinct splicing profile in primary T cells, particularly in activated T cells.

Finally, we evaluated the distribution of AS type (alternative 5’ or 3’ splice site usage, exon skipping and intron retention) observed in PRMT5 deficient activated T cells. Among the 2590 alternative junctions modulated by PRMT5, exon skipping (46.5%) was the most frequent with PRMT5 loss, closely followed by intron retention (28%), while alternative 5’ or 3’ junctions were less frequent (**Fig. 1E**). These results suggest that PRMT5 expression in T cells results in substantial and non-random changes in AS, which presumably modulate T cell biology and function. Our results raise the question of how PRMT5 is regulating splicing. Regulation could be via RNA binding protein (RBP) methylation, TCR signaling and/or RNA splicing protein methylation in T cells.

### PRMT5 methylates spliceosome Sm proteins and the RNA binding protein hnRNP K

PRMT5 has been described as a crucial player in spliceosomal assembly via symmetric dimethylation (SDM) of Sm proteins^23^. However, the spliceosomal or other methylation targets of PRMT5 in T cells are largely unknown. We hypothesized that PRMT5 methylates splicing proteins/regulatory factors in activated T cells. To study this, we first performed an unbiased pull down of SDM proteins in iCD4-PRMT5^fl/fl^ and iCD4-PRMT5^Δ/Δ^ T cells, using the SYM10 antibody that recognizes symmetrically dimethylated RGG and subjected the SDM target-enriched samples to mass spectrometry (**Fig. 2A**). We then used the Scaffold software to identify putative methylation targets. We focused on symmetrically dimethylated targets that were more highly recovered in activated iCD4-PRMT5^fl/fl^ condition compared to the activated iCD4-PRMT5^Δ/Δ^ condition. From this, we observed a number of splicing related proteins that were recovered in PRMT5-sufficient but not, or to a lesser extent, in PRMT5-deleted activated T cells. For our analysis, we graphed the raw spectral reads of the targets.

**Figure 2.**
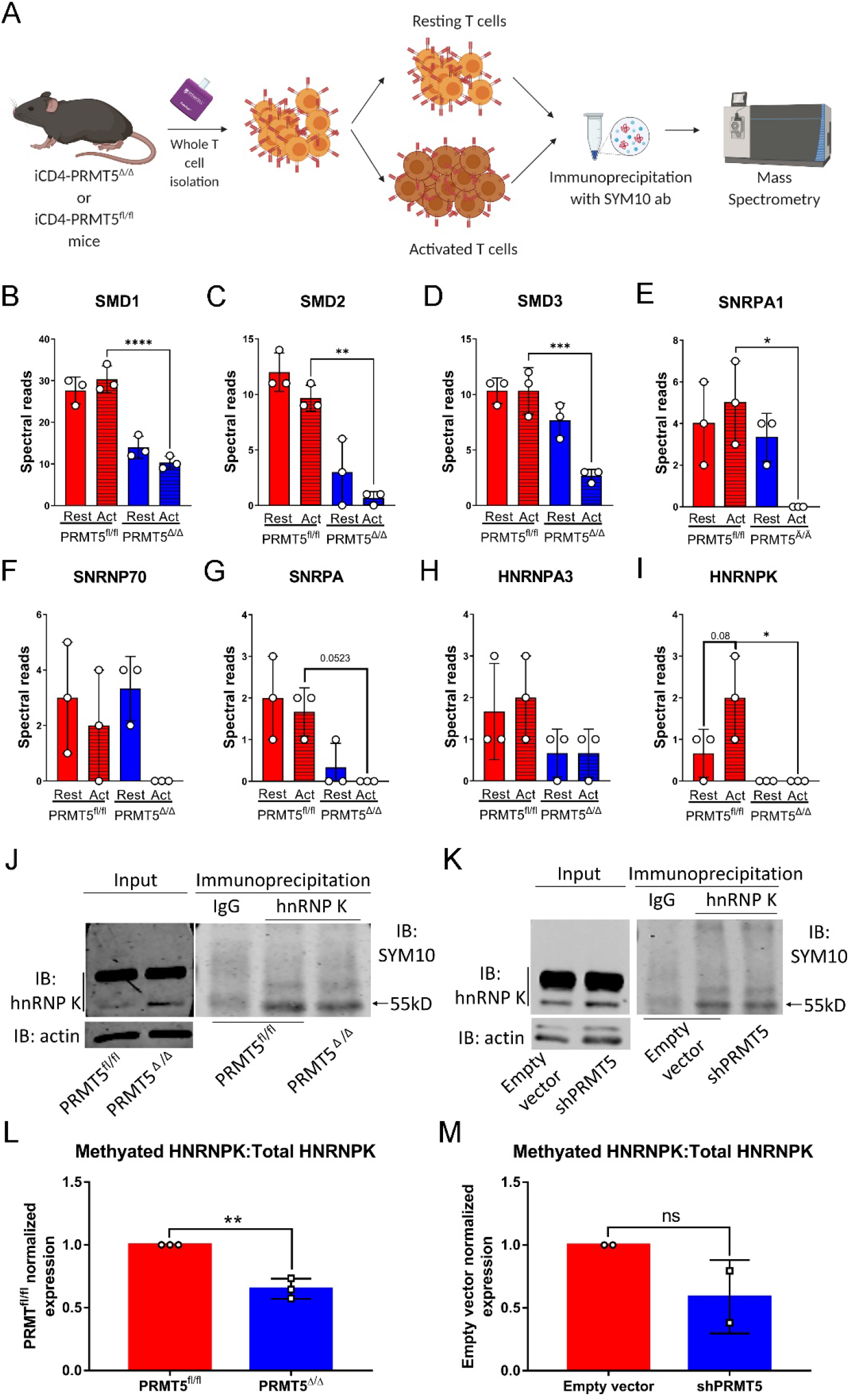
PRMT5 methylates spliceosome Sm proteins and the RNA binding protein hnRNP K. (**A**) Experimental design for Immunoprecipitation (IP)-Mass Spectrometry (MS) of 48hr activated, whole CD4^+^T cells negative selection from iCD4-PRMT5^fl/fl^ and iCD4-PRMT5^Δ/Δ^ mice. n=3 samples, combined from 9 age-matched mice in each respective group IP: pan-symmetric dimethylation antibody SYM10. MS: Liquid Chromatography with tandem mass spectrometry (LC-MS/MS). Figure created in Biorender. (**B – I**) SMD1 (**B**), SMD2 (**C**), SMD3 (**D**), SNRPA1 (**E**), SNRNP70 (**F**), SNRPA (**G**), HNRNPA3 (**H**) and HNRNPK (**I**) mass spectrometry spectral reads in protein lysates from resting and 48hr activated (act) whole CD4+ T cells from iCD4-PRMT5^fl/fl^ (**red**) and iCD4-PRMT5^Δ/Δ^ (**blue**) mice after IP with the pan-symmetric dimethylation antibody SYM10. One-way ANOVA followed by Sidak’s multiple-comparisons test was used. \**P* < 0.05, \*\**P* < 0.01, \*\*\**P* < 0.001, \*\*\*\**P* < 0.0001. Bar graphs display mean ± SD. (**J**) SYM10 immunoblot in 48hr activated CD4^+^ T cells from iCD4-PRMT5^fl/fl^ and iCD4-PRMT5^Δ/Δ^ mice after IP of HNRNPK. Band corresponding to HNRNP K is observed at 55kDa. Data representative of 3 independent experiments. (**K**) Jurkat cells with shRNA for PRMT5 were used in a hnRNP K IP and SYM10 IB. Empty vector is a scramble shRNA control. Band corresponding to hnRNP K is observed at 55kDa. Data representative of 2 independent experiments. (**L-M**) Quantification of HNRNPK IP bands for PRMT5^fl/fl^ and PRMT5^Δ/Δ^ normalized to HNRNPK from input in 48hr activated CD4^+^ T cells from iCD4-PRMT5^fl/fl^ and iCD4-PRMT5^Δ/Δ^ mice (**L**) or Jurkat T cells (**M**). Student’s *t* test. \*\**P* < 0.01. Bar graphs display mean ± SD.

Sm proteins SMD1, D2 and D3, which are responsible for the Sm-ring formation step required for spliceosome formation^24^ (**Fig 2 B-D**), were recovered significantly less in PRMT5 deficient activated T cells. However, methylated Sm protein recovery was similar between resting and activated wild-type T cells. Recovery of other splicing-related proteins such as SNRPA1, SNRNP70, SNRPA that help with spliceosomal assembly (aiding the binding of stem loop (SL)IV of U2 snRNA, SLI and SLII to U1 snRNA respectively)^25,26^ or HNRNPA3 which helps with cytoplasmic trafficking of RNA^27^ did not significantly decrease with PRMT5 deletion (**Fig. 2 E-H**). However, recovery of HNRNPK, an RBP that assists in the maturation of pre-mRNAs into mRNAs, stabilizes the mRNA during transport and controls the translation of the mRNA^28^ (**Fig. 2I**), followed the expected changes after T cell activation and PRMT5 loss. Specifically, HNRNPK recovery increased after T cell activation, when PRMT5 is induced, but decreased with PRMT5 deletion (**Fig. 2I**). Based on the role of hnRNP K in RNA splicing and the PRMT5-dependent recovery of methylated HNRNPK in activated T cells, we further validated this target in T cells. We performed a ‘reverse’ IP where we pulled down the target HNRNPK in activated T cells from iCD4-PRMT5^fl/fl^ and iCD4-PRMT5^Δ/Δ^ mice or control vs. PRMT5 shRNA-modified human Jurkat T cells and probed with SYM10 (**Fig. 2, KJ**). We observed significantly reduced detection of methylated HNRNPK in cells from PRMT5 knockout murine (**Fig. 2J, L**) and a trend in human (**Fig. 2K, M**) T cells, suggesting hnRNP K methylation contributes to PRMT5-dependent AS changes that occur upon T cell activation.

### T cell *Trpm4* exon inclusion is controlled by PRMT5

Our data so far support that PRMT5 promotes AS changes in activated T cells that have the potential to modulate T cell biology and function. To evaluate the immunological significance of genes whose alternative splicing is regulated by PRMT5, we ran the list through the immune effector processes node (GO: 0002697, **Fig. 3A**) in the gene ontology knowledgebase. Immune genes whose splicing is modulated by PRMT5 corresponded to subcategories in Fc receptor signaling (GO:0038093), TCR signaling (GO:0050852) and regulation of T cell cytokine production (GO:0002724). Within the regulation of T cell cytokine production subfamily, transient receptor potential melastatin 4 (Trpm4) was of interest in the context of our model because it has been shown to regulate Ca^2+^ signaling and IL-2 production^2^. Therefore, we studied PRMT5’s impact on *Trpm4* AS further.

**Figure 3.**
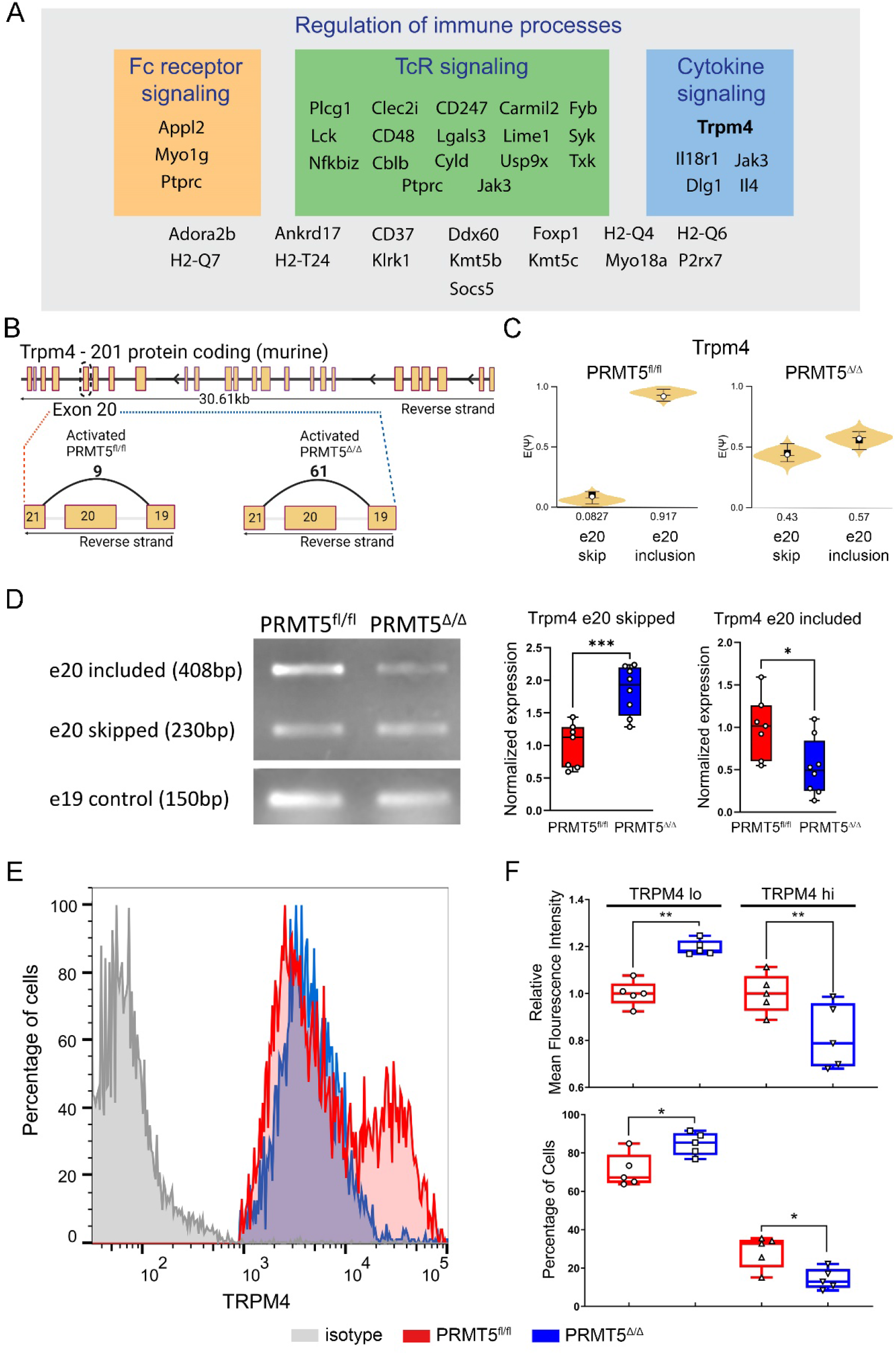
TRPM4 channel expression in T cells is affected by loss of PRMT5. (**A**) Regulation of immune processes is a key biological term discovered from gene ontology (GO) analysis. Within this umbrella we see an overlap with genes whose alternative splicing is modulated by PRMT5, belonging to Fc receptor signaling, TcR signaling and cytokine signaling. Grouping is based on the Panther classification system comparing against the *Mus musculus* reference genome. (**B**) Protein coding murine *Trpm4-201* splice map. Exon 20 skipping splice graphs for PRMT5^fl/fl^ and PRMT5^Δ/Δ^ 48 hr activated T cells are depicted. Numbers indicate the raw RNA-Seq reads for exon 20 skipping occurring in the two groups. Figure created in Biorender. (**C**) Violin plots from VOILA shows the expected Ψ value of *Trpm4* exon 20 skipping or inclusion in PRMT5^fl/fl^ act vs. PRMT5^Δ/Δ^ activated T cells. Ψ value is represented in a 0 to 1 range indicating the probability of the AS event occurring. (**D**) Agarose gel PCR amplifying exon 19-21, showing *Trpm4* exon 20 skipping in PRMT5^fl/fl^ and PRMT5^Δ/Δ^ T cells (middle band). A section of exon 19 was used as a control product (lower band). Quantification of semiquantitative PCR assaying the PRMT5^fl/fl^ normalized expression of exon 20 skipping or inclusion in PRMT5^fl/fl^ and PRMT5^Δ/Δ^ T cells. Student’s *t* test \*\*\*\**P* < 0.0001. Box-and-whisker plots (points = max to min, line = median, box = 25th–75th percentiles). Plots display mean ± SD. (**E**) Representative histogram overlay of TRPM4 staining in 48hr activated PRMT5^fl/fl^ and PRMT5^Δ/Δ^ T cells. PRMT5^fl/fl^ T cells are indicated in red and PRMT5^Δ/Δ^ T cells are in blue. Isotype control denoted in grey. (**F**) Relative mean fluorescence intensity (MFI) and percentage of cells depicting TRPM4 lo and TRPM4 hi peaks in PRMT5^fl/fl^ and PRMT5^Δ/Δ^ T cells. One way ANOVA, followed by Sidak’s multiple comparisons test. \**P* < 0.05, \*\**P* < 0.01. Box-and-whisker plots (points = max to min, line = median, box = 25th–75th percentiles). Plots display mean ± SD.

To visualize PRMT5-dependent LSV events in *Trpm4*, we used the VOILA tool within MAJIQ, which provides a sashimi plot that shows several exon junction connections entering or leaving a reference exon. This analysis showed that our PRMT5^Δ/Δ^ T cells have increased RNA-Seq reads for the skipping of exon 20 (61 vs. 9 in PRMT5^fl/fl^, **Fig. 3B**). This observation is better visualized in the percent spliced index (Ψ) provided by VOILA. The violin plots (**Fig. 3C**) show the inclusion or skipping probability of *Trpm4* exon 20 in the PRMT5^fl/fl^ and PRMT5^Δ/Δ^ conditions. We observe 91.7% usage of exon 20 inclusion in the PRMT5^fl/fl^ condition (**Fig. 3C, left**), which drops to 57% usage in the PRMT5^Δ/Δ^ condition (**Fig. 3C, right**). These *in silico* findings were confirmed in lab via semi-quantitative PCR amplification of a fragment encompassing exon 19 to exon 21. We observed a significant increase in the skipped product in the PRMT5 deficient condition and a significant decrease in the included/non-skipped product in the PRMT5 deficient condition (**Fig. 3D**), red corresponds to PRMT5 sufficient and blue to the PRMT5 deficient condition). To elucidate the biological significance of a loss of exon 20 in murine *Trpm4*, we consulted the Ensembl database. We found that the loss of exon 20 leads to nonsense mediated decay (NMD) due to an out-of-frame shift (178bp). Based on this, and the fact that there is an increase in the skipped product in the iCD4-PRMT5^Δ/Δ^ condition, we hypothesized that there is a loss of functional TRPM4 channels in the PRMT5 knockout T cells. We evaluated this by performing flow cytometry for TRPM4 in whole CD4 T cells. We observed that TRPM4^hi^ and TRPM4^lo^ populations can be observed in wild-type iCD4-PRMT5^fl/fl^ Th cells. However, loss of PRMT5 resulted in a significant loss of the Trmp4^hi^ population, and a significant increase in the TRPM4^lo^ population (**Fig. 3E-F**). Decreases in TRPM4 MFI were also observed for the TRPM4 hi population in PRMT5 deficient T cells (**Fig. 3F**). We interpret this result as lack of PRMT5 limiting the TRPM4^hi^ population in activated CD4 T cells, what likely impairs T cell activation and or expansion.

### PRMT5 promotes Calcium signaling, NFAT nuclear localization and IL-2 secretion

TcR signaling induces entry of Calcium (Ca^2+^), which acts as a secondary messenger in T cell signaling pathways^29^. To properly activate the transcriptional programs necessary for effective immune responses, appropriate Ca^2+^ signal amplitude and duration are necessary^30^, which requires depolarization via loss of other cations. TRPM4 is a Ca^2+^ activated Na^+^ channel that permits calcium oscillation by inducing depolarization, thereby allowing sustainably elevated Ca^2+^ levels^31^. Ca^2+^ in turn activates calcineurin and promotes NFAT nuclear translocation and *Il-2* gene transcription^32^. If PRMT5-dependent *Trpm4* AS alters TRPM4 function, we would expect altered Ca^2+^ signaling. To study whether PRMT5 modulates the calcium signaling profile in our mouse model, we performed a 600-second trace of calcium uptake in PRMT5^Δ/Δ^ and PRMT5^fl/fl^ T cells (**Fig. 4A**). Cells were kept in a zero-calcium media condition and treated with the ER calcium release inhibitor thapsigargin prior to addition of Ca^2+^ to the media. This strategy provides a system to specifically study cytosolic calcium entry and plasma membrane channel response. Quantification of the ‘plateau’ region of the trace after CaCl2 addition showed that the PRMT5^Δ/Δ^ T cells have a significant reduction in total cytosolic calcium uptake (**Fig. 4B**). To determine if the expected outcome of Ca^2+^ signaling in T cells, nuclear localization of NFAT transcription complexes, was also affected, we performed NFATc1 immunocytochemistry staining. We observed a decrease in nuclear localization in the PRMT5^Δ/Δ^ T cells (**Fig. 4C**, red - NFATc1, blue - DAPI). The quantification of these results confirmed a significant decrease in nuclear localization of NFATc1 in PRMT5^Δ/Δ^ T cells (**Fig.4D**). Finally, it’s been established that NFAT nuclear localization in activated T cells promotes the expression of interleukin (IL)-2 and we have previously observed decreased IL-2 after PRMT5 inhibition or KO. As expected from our prior work and the role of NFAT as an IL-2 driver, PRMT5 deleted T cells secreted far less IL-2 upon T cell activation (**Fig. 4E**).

**Figure 4.**
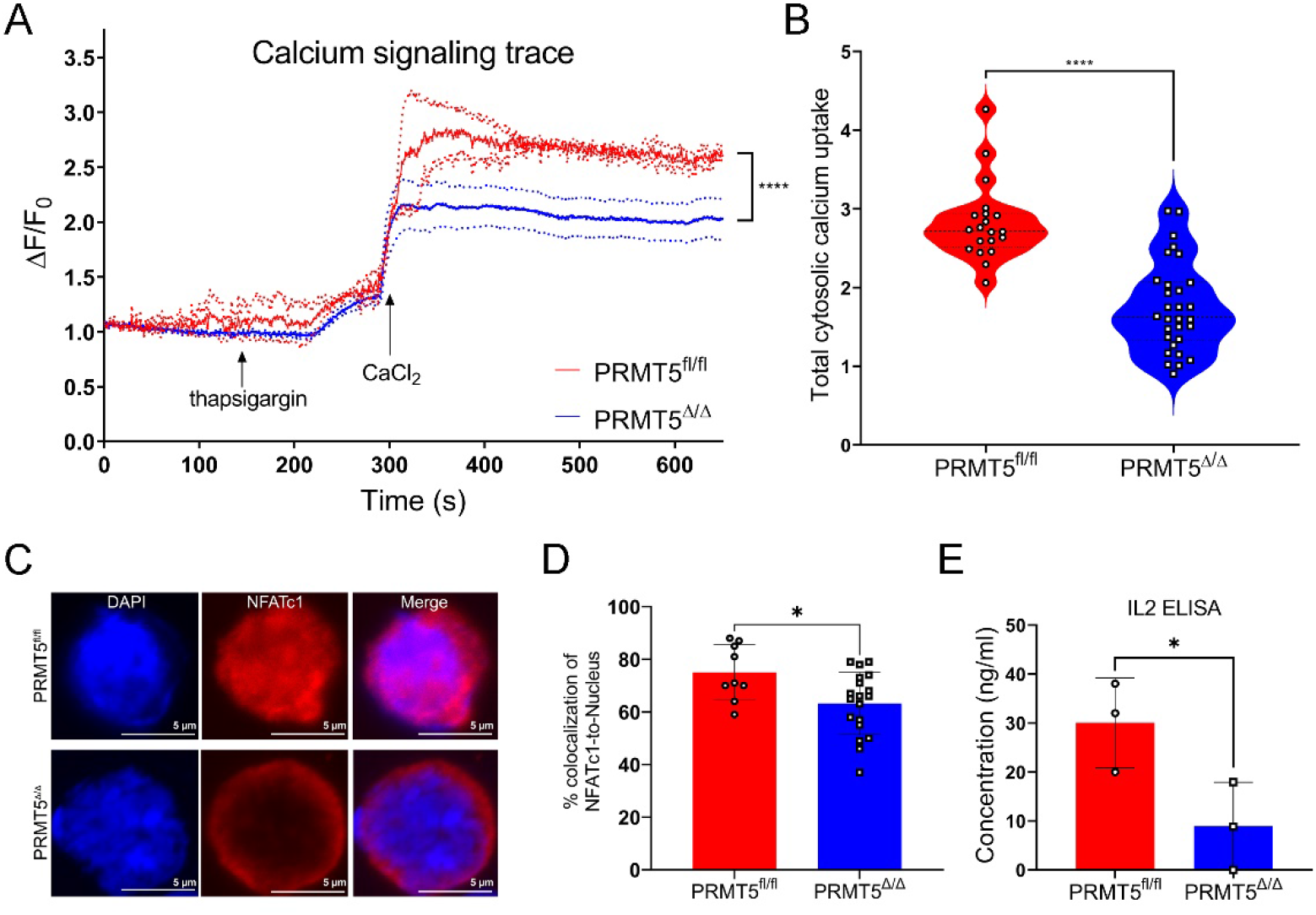
PRMT5 loss in T cells reduces calcium processivity and impacts IL-2 production. (**A**) Live cell Fluo-4 trace studying total cytosolic calcium uptake in PRMT5^fl/fl^ and PRMT5^Δ/Δ^ T cells. PRMT5^Δ/Δ^ T cells show a lower threshold of calcium uptake. Trace is represented as mean ± SEM, with dashed lines representing the SEM (**B**) Violin plot showing total cytosolic calcium uptake. Data points for quantification were selected from the plateau region of the trace spike after CaCl2 addition. n= 20 time points in PRMT5^fl/fl^ condition, and n= 30 time points in PRMT5^Δ/Δ^ condition. Student’s *t* test \*\*\*\**P* < 0.0001. Plot is displayed as mean ± SEM. Points = max to min, line = median. (**C**) NFAT localization in PRMT5^fl/fl^ and PRMT5^Δ/Δ^ T cells. Clear nuclear translocation is observed in PRMT5^fl/fl^ samples. Colocalization of NFAT (red) and DAPI (blue) is used to determine nuclear translocation. (**D**) Quantification of percent colocalization of NFATc1-to-Nucleus in PRMT5fl/fl and PRMT5^Δ/Δ^ T cells activated T cells, as in (**C**). Quantification of individual cells have been plotted and mean and SEM were used to plot the comprehensive data. Student’s *t* test. (**E**) Quantification of IL-2 ELISA. ELISA performed on supernatants from 48 hr activated PRMT5^fl/fl^ and PRMT5^Δ/Δ^ T cells. Student’s *t* test \**P* < 0.05. Plots are displayed as mean ± SD. Points = max to min.

## Discussion

The main goal of this work was to explore the role of PRMT5 on AS changes in T cells and identify methylation targets of PRMT5 in T cells. We find that PRMT5 symmetrically dimethylates several proteins involved in RNA processing, including SmDs and HNRNPK, and is required for a portion of the AS pattern induced by T cell activation. We additionally found that PRMT5 modulates the splicing of a sodium channel that modulates calcium processivity, namely *Trpm4*, and promotes NFAT signaling and IL-2 production in Th cells.

A substantial contribution of alternative splicing to gene expression changes induced by T cell activation was initially recognized in 2007^13,15^. More recently, it has been shown that a number of AS changes in activated T cells translate to differential protein isoform expression and changes in T cell function^16,33,34^. T cell activation-dependent AS changes impact genes that modulate a range of T cell processes, from signaling, migration or fate decisions, to proliferation ^15,19,35^. We have previously shown that PRMT5 induction after T cell activation^4^ promotes activation-induced cell cycle progression^12^ and proliferation^2,3^. The MAJIQ analyses of PRMT5 sufficient and deficient T cells in the current work show that PRMT5 controls approximately 16% of T cell activation induced AS shifts. Such shifts occurred in genes involved in TcR, Fc and cytokine signaling, as well as other immune processes. In addition, we provide evidence that control of AS by PRMT5 is active in primary murine T cells in which loss of PRMT5 impacts T cell proliferation, NFAT signaling and IL-2 secretion. While our work does correlatively link these processes, future work will need to demonstrate whether specific AS changes in specific genes are necessary and/or sufficient to influence function.

The contribution of PRMT5 to alternative splicing was first recognized in plants^36^. Since then, the role of PRMT5 in splicing has been studied in hematopoietic stem cells^37^, neural stem/progenitor cells^38^, monocytic leukemia cells^39^ and murine glioma cells^40^, among others. These studies evidenced that PRMT5 mediated splicing is crucial in modulating DNA repair genes^37^, MDM4^38^ and avoiding intron retention^39,40^. Metz *et al*^41^ studied the impact of PRMT5 small molecule inhibitors in human T cell splicing and concluded that a global reduction in SDM levels altered the splicing of a limited set of mRNA transcripts and selectively prevented TcR and pattern recognition receptor (PRR) dependent upregulation of *IFNB1* and *IFNL1.* We now show the extent to which genetic loss of PRMT5 controls splicing and find that PRMT5 controls a substantial portion of TcR-induced AS changes. As Metz et al found, not all mRNAs in our dataset appear to require PRMT5 for processing. How exactly this is achieved is currently unknown. However, TcR-induced splicing has been shown to be highly dependent on CELF2 induction and binding to specific mRNA sites^17,35,42^, leading to the intriguing possibility that interactions between PRMT5 and CELF2 may contribute to selective splicing regulation of a group of transcripts required for activated T cell function.

We observe significantly less SmD1, SmD2, SmD3 and SNRPA1 methylation in activated PRMT5 deficient T cells. Sm proteins were the first identified methylation targets of PRMT5 that modulate RNA processing. Specifically, SmD1, SmD3, SmB and SmB’ were found to be SDM on RG motifs by PRMT5. Methylated Sm proteins then bind SMN and accelerate U snRNP assembly^23,43–45^. Therefore, PRMT5 appears to regulate early stages of spliceosomal assembly, during SMN binding and Sm ring formation. We also observe a significant loss of SDM of hnRNP K with PRMT5 loss in mouse activated T cells. hnRNPs are involved in RNA metabolism processes such as mRNA export, localization, stability and translation^46^. hnRNPA1 methylation by PRMT5 promotes IRES-dependent translation of CCND1, MYC, HIF1a and ESR1 genes^47^. Additional work is now showing that hnRNPs modulate alternative splicing of pyruvate-kinase isozyme splicing^48^, insulin receptor gene splicing^49^, CD45 alternative splicing^50^ and regulate innate immunity gene control in macrophages^51^. Although further work demonstrating that methylated hnRNP mediates the observed AS changes will be necessary, our data suggest a role for hnRNP K methylation in PRMT5-dependent AS changes observed in T cells.

We found and validated *Trpm4* as an alternative splicing target of PRMT5 that is, as a consequence, substantially repressed at the protein level in PRMT5 deficient activated T cells. The TRPM family of channels is expressed in several immune cells^52^, where it controls cell proliferation, survival and cytokine production^53^. While research on how TRPM4 contributes to T cell Ca^2+^ release, NFAT signaling and IL-2 secretion has yielded contradictory results^31,53^, we observe reductions of all three parameters in PRMT5 ^Δ/Δ^ T cells. We hypothesize this is due to reduced TRPM4 leading to lower calcium processivity. This finding could be important to explore when targeting ion channels to treat autoimmune neuroinflammation. Given the fact that the lower levels of calcium lead to reduced NFATc1 nuclear localization, it is exciting to consider if PRMT5 inhibition could be an efficient approach to modulating overactive T cell subsets.

In summary, our work shows that PRMT5 is an important mediator of TcR-induced AS in T cells and suggest that altered methylation in splicing proteins and changes in Ca^2+^/NFAT signaling underlie TcR expansion defects in PRMT5 deficient T cells. Additional studies will be needed to conclusively demonstrate the contribution of specific PRMT5-dependent AS changes to concrete T cell and disease phenotypes. This work and future studies may guide development of drugs targeting these processes and provide benefit to patients with autoimmune and other T cell mediated diseases.

## Supporting information

Table for Mass spectrometry targets

Supplemental fig. - Resting iCD4-PRMT5 fl/fl vs iCD4-ORMT5 D/D

## Acknowledgements

This work was supported by funds from the NIH National Institute of Allergy and Infectious Diseases grants R01AI121405 and 1R21AI127354 (both to MGA), The Ohio State University School of Health and Rehabilitation Sciences start-up funds (to MGA), the Comprehensive Cancer Center Mass Spectrometry Resource (Core Cancer Center Support Grant P30CA016058), the NIH National Cancer Institute grant 01-CA186729 (to PNT), the Pelotonia Postdoctoral Fellowship (to GL), and the Center for Clinical and Translational Science (CCTS) Award Number Grant UL1TR002733 from the National Center For Advancing Translational Sciences. The content is solely the responsibility of the authors and does not necessarily represent the official views of the National Center For Advancing Translational Sciences or the NIH. We would like to thank the Genomic Services Laboratory of the Abigail Wexner Research Institute at Nationwide Children’s Hospital for their help with RNA-Seq. We thank Amy Wetzel, Shireen Woodiga, Anthony Miller, and Saranga Wijeratne of the Genomic Services Laboratory at the Abigail Wexner Research Institute at Nationwide Children’s Hospital, Columbus, Ohio, for their help with sample QC, library preparation, RNA-Seq, and analysis of data. We would also like to thank Liwen Zhang and Sophie Harvey from the genomics shared resource at OSU for their help with mass spectrometry and analysis of data.

