## Supplemental fig. - Resting iCD4-PRMT5 fl/fl vs iCD4-ORMT5 D/D for "PRMT5 promotes symmetric dimethylation of RNA processing proteins and modulates activated T cell alternative splicing and Ca^2+^/NFAT signaling"

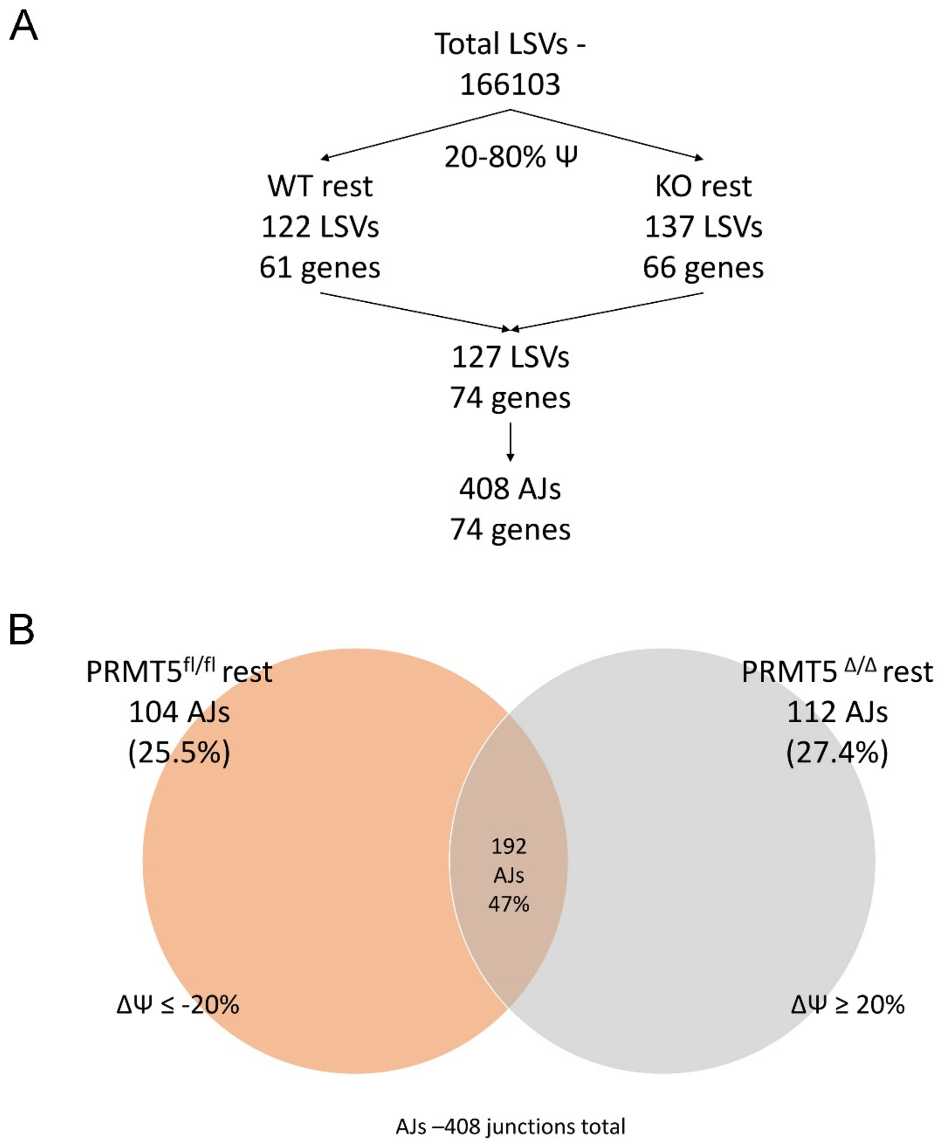


Supplemental Fig 1. MAJIQ analysis of resting iCD4-PRMT5$^{fl/fl}$ and iCD4-PRMT5$^{\Delta/\Delta}$ T cells

(**A**) MAJIQ workflow identifying significantly utilized AJs by comparing LSVs in resting iCD4-PRMT5$^{fl/fl}$ vs. iCD4-PRMT5$^{\Delta/\Delta}$ T cells. LSVs with at least two or more exon junctions within the 20-80% percent spliced index (Ψ) of reads were calculated. (**B**) Allocation of LSVs with two or more AJs in iCD4-PRMT5$^{fl/fl}$ resting and iCD4-PRMT5$^{\Delta/\Delta}$ resting groups. Shift in AJ usage is denoted as the difference in Ψ (ΔΨ) and is set at a minimum of 20% between conditions.
